# Small RNA Mcr11 regulates genes involved in the central metabolism of *Mycobacterium tuberculosis* and requires 3’ sequence along with the transcription factor AbmR for stable expression

**DOI:** 10.1101/616912

**Authors:** Roxie C. Girardin, Kathleen A. McDonough

## Abstract

*Mycobacterium tuberculosis* (Mtb), the etiologic agent of tuberculosis, must adapt to host-associated environments during infection by modulating gene expression. Small regulatory RNAs (sRNAs) are key regulators of bacterial gene expression, but their roles in Mtb are not well understood. Here, we address the expression and function of the Mtb sRNA Mcr11, which is associated with slow bacterial growth and latent infections in mice. We found, by using biochemical and genetic approaches, that the AbmR transcription factor and an extended region of native sequence 3’ to the *mcr11* gene enhance production of mature Mcr11. Additionally, we found that expression of Mcr11 was unstable in the saprophyte *Mycobacterium smegmatis*, which lacks an *mcr11* orthologue. Bioinformatic analyses used to predict regulatory targets of Mcr11 identified 9-11 nucleotide regions immediately upstream of Rv3282 and *lipB* with potential for direct base-pairing with Mcr11. *mcr11*-dependent regulation of Rv3282, *lipB,* Rv2216 and *pknA* was demonstrated using qRT-PCR in wild type versus *mcr11*-deleted Mtb and found to be responsive to the presence of fatty acids. These studies establish that Mcr11 has roles in regulating growth and central metabolism in Mtb that warrant further investigation. In addition, our finding that multiple factors are required for production of stable, mature Mcr11 emphasizes the need to study mechanisms of sRNA expression and stability in TB complex mycobacteria to understand their roles in TB pathogenesis.

**Author Summary:** Bacterial pathogens must continuously modulate their gene expression in response to changing conditions to successfully infect and survive within their hosts. Transcription factors are well known regulators of gene expression, but there is growing recognition that small RNAs (sRNAs) also have critically important roles in bacterial gene regulation. Many sRNAs have been identified in *M. tuberculosis* (Mtb), but little is known about their expression, regulatory targets or roles in Mtb biology. In this study, we found that the Mtb sRNA Mcr11, which is expressed at high levels in slowly replicating Mtb and during mouse infection, regulates expression of several target genes involved in central metabolism. Importantly, we also discovered that *mcr11* has unexpected requirements for stable expression in mycobacteria. In particular, we identified RNA sequence elements immediately downstream of *mcr11* that enhance transcription termination and production of mature Mcr11 RNA in TB-complex mycobacteria. Meanwhile, ectopic expression of Mcr11 was unstable in a non-pathogenic strain of mycobacteria, suggesting that factors specific to pathogenic mycobacteria are required for the stable production of Mcr11. These studies identify sRNA stability as a new frontier for understanding gene expression in Mtb.

## Introduction

Tuberculosis (TB) remains a major threat to global public health, with at least 10 million new, incident cases and 1.3 million deaths due to TB in 2017 (1). The basic biology of *Mycobacterium tuberculosis* (Mtb), the etiological agent of TB, is poorly understood despite its importance to the development of new therapeutic interventions. Mtb can adopt a specialized physiological state within host tissues, which renders the bacteria phenotypically drug resistant and viable despite extended periods of slow or non-replicating persistence (NRP) (2). NRP and phenotypic drug resistance pose particular challenges for intervention, making it critical to understand the regulatory processes that enable Mtb to adapt to host conditions.

Bacterial and host factors that contribute to NRP and slow growth are still being defined (3). Host-associated environmental cues that result in metabolic remodeling and a shift away from active growth toward a state of persistence include hypoxia, nitrosative stress, redox stress, nutrient starvation, as well as adaptation to cholesterol and fatty acid metabolism (4-9). Isocitrate lysases (ICLs) are used by bacteria to maintain growth on fatty acids through the glyoxylate shunt, and are critical for Mtb’s ability to survive on fatty acids and within the host (10-12). Surprisingly, ICLs are also required for growth on carbohydrate carbon sources (13), and gluconeogenesis is required for Mtb virulence (14-16). Additionally, lipoylated enzymes involved in the citric acid cycle, such as lipoamide dehydrogenase (Lpd) and dihydrolipoamide acyltransferase (DlaT), are necessary for Mtb survival in the host and viability during NRP (17-19). However, factors that regulate these processes are not well understood.

Gene expression studies have provided critical insights into the regulation and function of proteins like transcription factors that modulate gene expression as Mtb adapts to the host environment during infection (20, 21). The additional role of sRNAs in gene regulation is recognized in other bacteria (22), and several sRNAs whose expression is responsive to stress and/ or growth phase have been identified in mycobacteria (23-28). (23-26, 29-31). It also has been observed that over expression of some sRNAs leads to slow or delayed growth in mycobacteria (23, 32). However, the importance of these differentially expressed sRNAs to the adaptation and survival of Mtb during periods of stress has not been fully addressed.

The intergenic sRNA Mcr11 (*ncRv11264Ac*) is one of the best-studied mycboacterial sRNAs (24, 26, 29, 33). Expression of Mcr11 is regulated in response to advanced growth phase and levels of the universal second messenger 3’,5’-cyclic adenosine monophosphate (cAMP) (24, 26, 29). Additionally, Mcr11 is abundantly expressed by Mtb in the lungs of chronically infected mice (26) as well as in hypoxic and non-replicating Mtb (32). Transcriptional regulators of *mcr11* expression include the product of the adjacent, divergently expressed gene *abmR* (*Rv1265)* (33), and the cAMP binding transcription factor (TF) CRP_MT_ (29). The structure and function of Mcr11 in TB-complex mycobacteria is unknown, prompting us to further characterize this sRNA.

Here, we report that cis-acting, extended, native sequence 3’ to Mcr11 enhances production of Mcr11 in mycobacteria. Optimal Mcr11 termination efficiency needed the transcription factor AbmR and was regulated by growth phase in Mtb and *Mycobacterium bovis* BCG, but not in the fast-growing saprophyte *Mycobacterium smegmatis*. We found that *mcr11* regulates expression of *lipB* and *Rv3282,* which contribute to central metabolic pathways associated with NRP and slow growth in Mtb. In addition, Mcr11 affected growth of both Mtb and BCG in hypoxic conditions without fatty acids. This study identifies TB complex-specific cis and trans factors required for stable Mcr11 expression while providing the first report of *mcr11*-dependent regulation of gene targets in Mtb.

## Results

### Modeling secondary structure of Mcr11

Previous studies established two 5’ ends of Mcr11 at chromosomal positions 1413224 and 1413227 in *Mycobacterium tuberculosis* H37Rv (24, 29), but the 3’ end of Mcr11 is poorly defined. Preliminary efforts to express Mcr11 based on size estimates from prior Northern blot experiments were not successful, despite the well-mapped 5’ end of the sRNA. Secondary structural features of sRNAs greatly contribute to their function, and RNA folding is most strongly influenced by the nucleotide sequence of the RNA (34, 35). We reasoned that defining the precise boundaries of Mcr11 could help in identifying its function.

We mapped the 3’ end of Mcr11 to chromosomal positions 1413107 and 1413108 in *Mycobacterium tuberculosis* (Mtb) using 3’ rapid amplification of cNDA ends (RACE) and Sanger sequencing (Figure 1A). These 3’ ends are 120 and 119 nucleotides (nt) downstream from the most abundant previously mapped 5’ end at position 1413227 (24). Our mapped 3’ ends vary 3-4 nucleotides from the 3’ end at chromosomal position 1413111 inferred by deep sequencing (36), and are 13-14 nt shorter than the 3’ end estimated by cloning (24) (Figure 1A). These results indicate that Mcr11 is a transcript between 117 and 121 nt long, which is consistent with its observed size by Northern blot (24, 26, 29).

**Figure 1.**
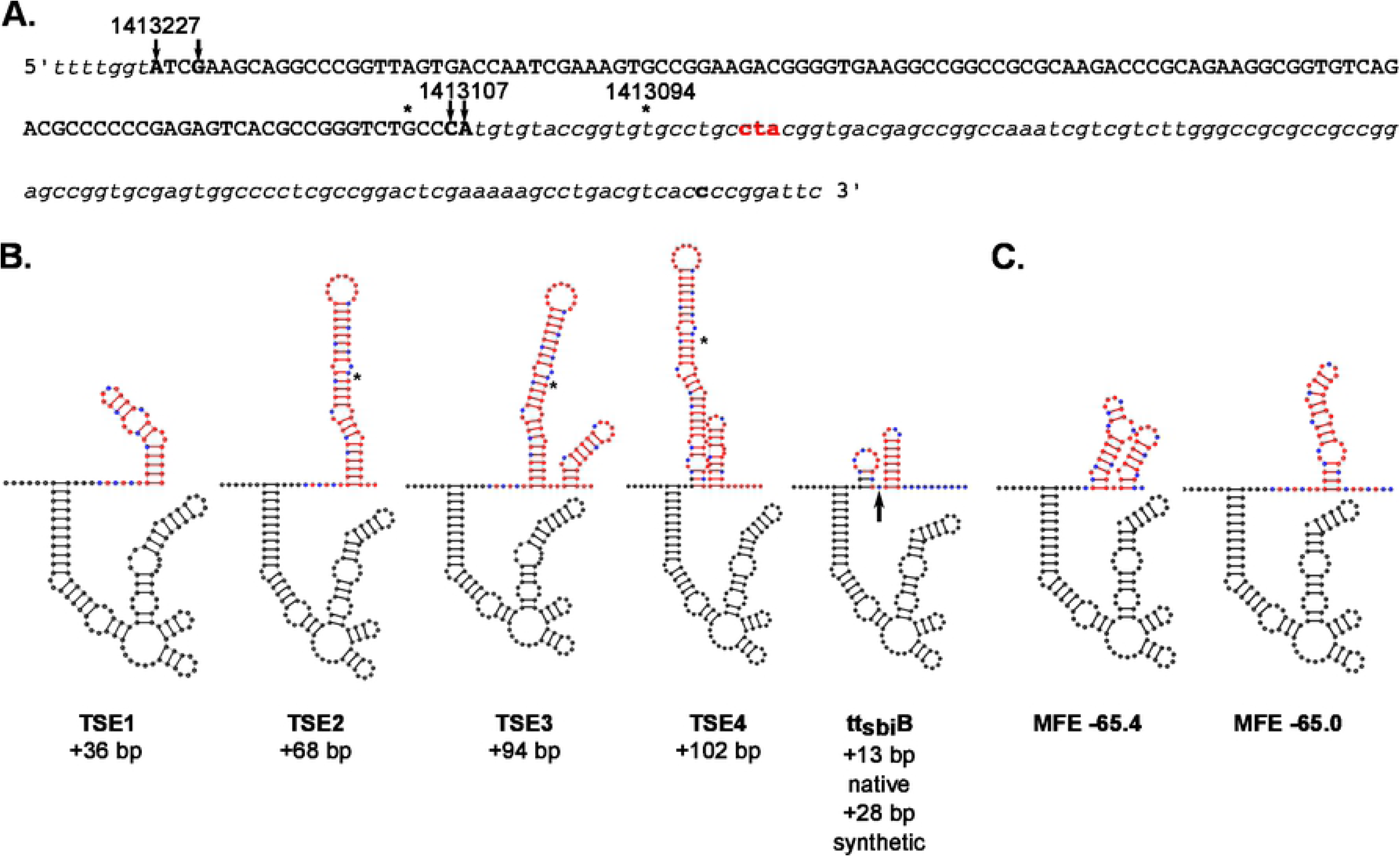
Secondary Structure modeling of 3’ sequences beyond the mapped 3’ end of Mcr11. A. The DNA sequence of the *mcr11* gene and extended 3’ sequence used for modeling experiments is shown. The *mcr11* gene is shown in capital letters, and nucleotides where 5’ and 3’ boundaries have been mapped by RACE are indicated by arrows and shown in capitalized, bolded black text. Flanking sequences are in italics, and the Rv1264 stop codon in bolded in red text. The last nucleotide on the 3’ side of mcr11 that was included in modeling experiments is in bold, lowercase black text. An asterisk indicates 3’ ends reported by DiChiara et al and DeJesus et al. Positions of mapped nucleotides on the Mtb chromosome are shown above the text. B. Secondary structure diagrams of Mcr11 from 5’ position 1413227 and extended 3’ native sequences. The synthetic idealized intrinsic termination control (tt_sbi_B) is also modeled onto Mcr11 from the longest 3’ end reported at position 1413094; the last nucleotide of native sequence 3’ to Mcr11 is indicated by a black arrow. Black nucleotides indicate bases in the mapped boundaries of the *mcr11* gene, red base pairs indicate base pairs beyond the mapped 3’ end of Mcr11 at position 1413107, except all uracils (Us) are shown in blue. An asterisk indicates that last nucleotide modeled in (C.). C. The secondary structure models with the lowest minimum free energies (MFE) of a shorter untested native Mcr11 TSE with the most U rich sequence at the 3’ end of the putative terminator. The MFE of each structure is shown below each diagram.

The mapped 3’ ends of Mcr11 at chromosomal positions 1413107/ 1413108 varied by 3-13 nt from previous estimates of the 3’ end of Mcr11, so we considered the possibility that this variance could be functionally significant. Transcriptional termination occurs in prokaryotes by two described mechanisms: Rho-dependent termination and intrinsic, Rho-independent termination (37-39). Rho-dependent termination requires the association of the ATP-hydrolyzing molecular motor Rho with the nascent RNA, which is typically cytosine-enriched, guanosine-depleted and unstructured at the 3’ end (40). Intrinsic, Rho-independent termination is governed by the formation of a structured termination hairpin that is usually followed a poly-U tail, and is highly dependent upon the sequence of the nascent RNA. Several mycobacterial sRNAs have multiple reported 3’ ends, suggestive of Rho-dependent termination and/or post-transcriptional processing (23, 30, 31). However, relatively little is known about transcriptional termination in Mtb, particularly at sRNAs (41-45).

We modeled the secondary structure of Mcr11 with varying lengths of extended, native 3’ RNA sequence by using RNA Structure (46) to reveal potential functional features. Mcr11 was predicted to be highly structured, with few single-stranded (ss) regions available for potential base-pairing with regulatory target RNA sequences (Figure 1B). Use of either mapped 5’ end of *mcr11* produced the same Mcr11 secondary structure, as these 5′ nucleotides are predicted to be unpaired (Figure 1B). Modeled structures of extended native sequence 3’ to our mapped ends of *mcr11* revealed additional highly structured motifs with similarity to predicted Rho-independent terminators (RITs) downstream of *mcr11* in mycobacteria (43, 44). However, none included the long, characteristic poly-U tail found in RITs of other bacteria (38). A 10 nt region containing a sub-optimal poly-U tract was identified by scanning sequence downstream of the mapped 3’ end of Mcr11 in the constructs that contained 3’ sequence elements (TSEs) 2-4 (Figure 1A). Modeling the secondary structure of this sequence revealed a stem loop structure with a 7nt trail containing 4 discontinuous Us (Figure 1C).

### Measurement of TSE function in Msm

We tested the impact of different *mcr11* TSEs on the transcriptional termination of Mcr11 using promoter:*mcr11:TSE:GFPv* reporter constructs (Figure 2A-B). We reasoned that transcriptional termination levels could be inferred by using GFPv fluorescence as a relative measure of transcriptional read-through beyond the *mcr11* gene (Figure 2A). The relative percentage of termination for each TSE was calculated by dividing the observed fluorescence of each P*mcr11:TSE:GFPv* strain by the observed fluorescence of an *mcr11* promoter-only:*GFPv* construct (P*mcr11:GFPv*), subtracting the product from 1, and multiplying the result by 100. Promoterless *GFPv* served as a negative assay control, and an *mcr11*-independent promoter:*GFPv* fusion construct served as a positive assay control. A positive intrinsic termination control was generated by fusing *mcr11* and a small amount of native 3’ trailing sequence to a synthetic RIT (tt_sbi_B) that has been shown to function at ~98% efficiency with Mtb RNAP *in vitro* (Figure 1B) (42).

**Figure 2.**
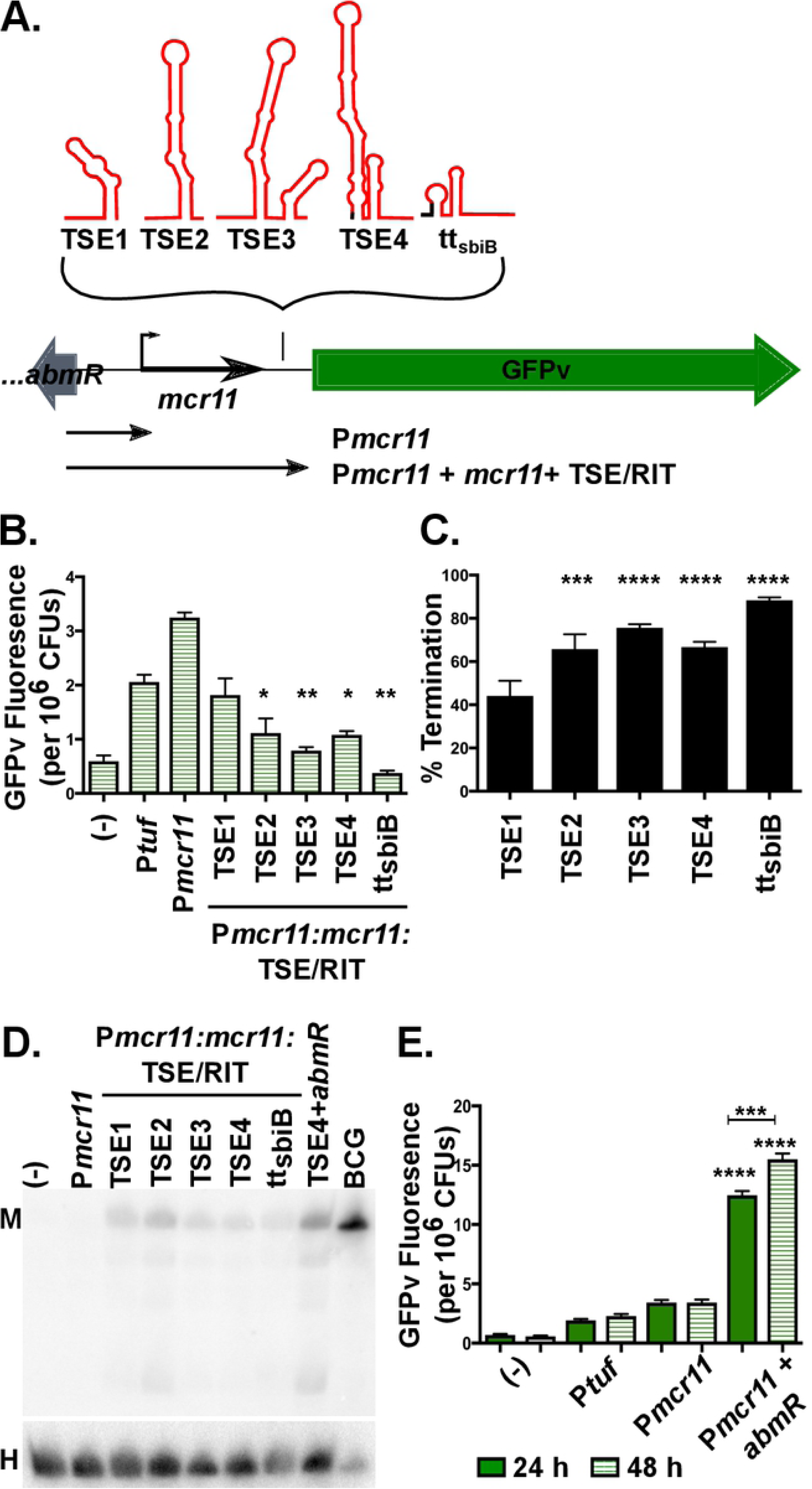
TSEs decrease transcriptional read-through of Mcr11-GFPv reporters, but Mcr11 is not robustly expressed in Msm, even in the presence of *abmR* from Mtb. A. A schematic representation of GFPv fluorescence-based reporter assay to determine the functionality of TSEs, and a synthetic idealized intrinsic termination control (tt_sbi_B). B. GFPv fluorescence assay used to measure promoter activity and read-through of TSEs of *mcr11* in late stationary phase Msm, which lacks a native *mcr11* locus. The promoter P*tuf* served as a positive control. The various TSE constructs tested are indicated underneath the corresponding bar. Statistical comparisons are relative to P*mcr11*. C. The % termination of constructs tested in (B.). Statistical comparisons are relative to TSE1. D. Northern blot analysis of Mcr11 expression (M) in Msm with various *mcr11* + TSE constructs. Ten ug of total Msm RNA was loaded, whereas 3 ug of the BCG positive control was loaded. The HisT tRNA (H) was used as a loading control. E. GFPv fluorescence assay used to measure promoter activity in Msm. Statistical comparisons are relative to P*mcr11* or between 24h and 28h as indicated. Results are the means of 3 biological replicates. Statistical analysis conducted with an unpaired, 2-tailed Student’s t-test. Asterisks indicate significance as follows: * p<0.05, ** p<0.01, *** p<0.001, **** p<0.0001.

Our initial termination experiments measuring TSE efficiency were performed in Msm, because it lacks an *mcr11* orthologue and thus has no basal Mcr11 expression. The inclusion of any TSE sequence at the 3’ end of *mcr11* resulted in decreased transcriptional read-through of the *mcr11*:GFPv fusion reporter, as compared to the relative levels of GFPv fluorescence for P*mcr11* alone (Figure 2B). This result suggested that all TSEs supported transcriptional termination to varying degrees. Longer TSEs had greater termination efficiency, but the positive tt_sbi_B intrinsic termination control exhibited the strongest mean termination efficiency at 88% (Figure 2C). Paraformaldehyde-fixed duplicates of each sample were subjected to flow cytometry analysis to determine if the mean GFPv fluorescence observed in our plate-based assay was reflective of the fluorescence observed in individual cells, or if there were populations of cells with varying levels of fluorescence. A single, homogenous population of fluorescent cells was observed for each reporter, demonstrating that mean fluorescence measured in the plate reader assay was representative of the fluorescence in individual cells (Supplemental Figure 2). From these data, we concluded that mean fluorescence reflected relative termination efficiency for each TSE reporter.

Northern blot analysis was performed to further evaluate the expression of Mcr11 in Msm. We noted that the size of Mcr11 did not vary in Msm despite the presence of TSEs with various lengths, suggesting that TSEs are rapidly processed off of the mature sRNA if they are transcribed (Figure 2D). Mcr11 was also present at very low levels in Msm when compared to a BCG control for which 1/3 the amount of total RNA present in the Msm samples was loaded in the gel (Figure 2D). These results are consistent with the possibility of reduced Mcr11 expression and/or stability in Msm compared to BCG.

### TB complex-specific factors are required for stable Mcr11 expression

*mcr11* expression varies across growth phases in TB complex mycobacteria (24, 33), so we examined the effect of growth phase on *mcr11* expression in Msm. Fluorescence from the GFPv transcriptional reporters was measured at mid-log and late stationary phase. Surprisingly, we observed that P*mcr11* activity was not increased by advancing growth phase when expressed in Msm (Figure 2E), suggesting that a positive regulatory factor was absent.

Previously, we showed that the divergently expressed DNA-binding protein AbmR is a growth phase-responsive activator of *mcr11* expression (33). Although the Msm orthologue (*MSMEG_5010)* of *abmR* displays both high amino acid sequence identity (68.75%) and amino acid similarity (83.51%) to AbmR (33), we hypothesized that *abmR* has functionality that the Msm orthologue lacks. Thus, we added a copy of the Mtb *abmR* locus upstream of the P*mcr11*::*GFPv* fusion sequences and tested the activity of P*mcr11* in response to growth phase in Msm. The addition of *abmR* significantly increased P*mcr11* activity in Msm, and rendered P*mcr11* activity responsive to growth phase (Figure 2E), as previously observed in BCG and Mtb (33). However, inclusion of *abmR* did not significantly increase the amount of stable Mcr11 detected by Northern blot in Msm (Figure 2D), indicating that Mcr11 is unstable when transcribed in Msm. Together, these results show that Mtb *abmR* regulates *mcr11* expression at the transcriptional level, but is not sufficient for stable Mcr11 expression in a non-native environment.

Further studies in TB-complex mycobacteria showed that the termination efficiencies of *mcr11* TSEs in mid-log phase BCG were similar to those observed in Msm (Figure 3), but only BCG showed enhanced *mcr11* TSE termination efficiencies in late stationary phase. This growth-phase dependent increase in termination efficiency in BCG was strongest for TSE1, which improved from approximately 40% to 70% (Figure 3C). Overall termination efficiencies of *mcr11* TSEs were also greater in virulent Mtb (H37Rv) than in BCG or Msm (Figure 3D-F). Deletion of *abmR* caused a small but significant decrease in *mcr11* termination efficiencies from all constructs, with the TSE1 short hairpin being the most strongly affected (Supplemental Figure 3). Together, these data indicate that multiple trans-acting factors specific to the genetic background of TB-complex mycobacteria facilitate the transcriptional termination and production of stable Mcr11.

**Figure 3.**
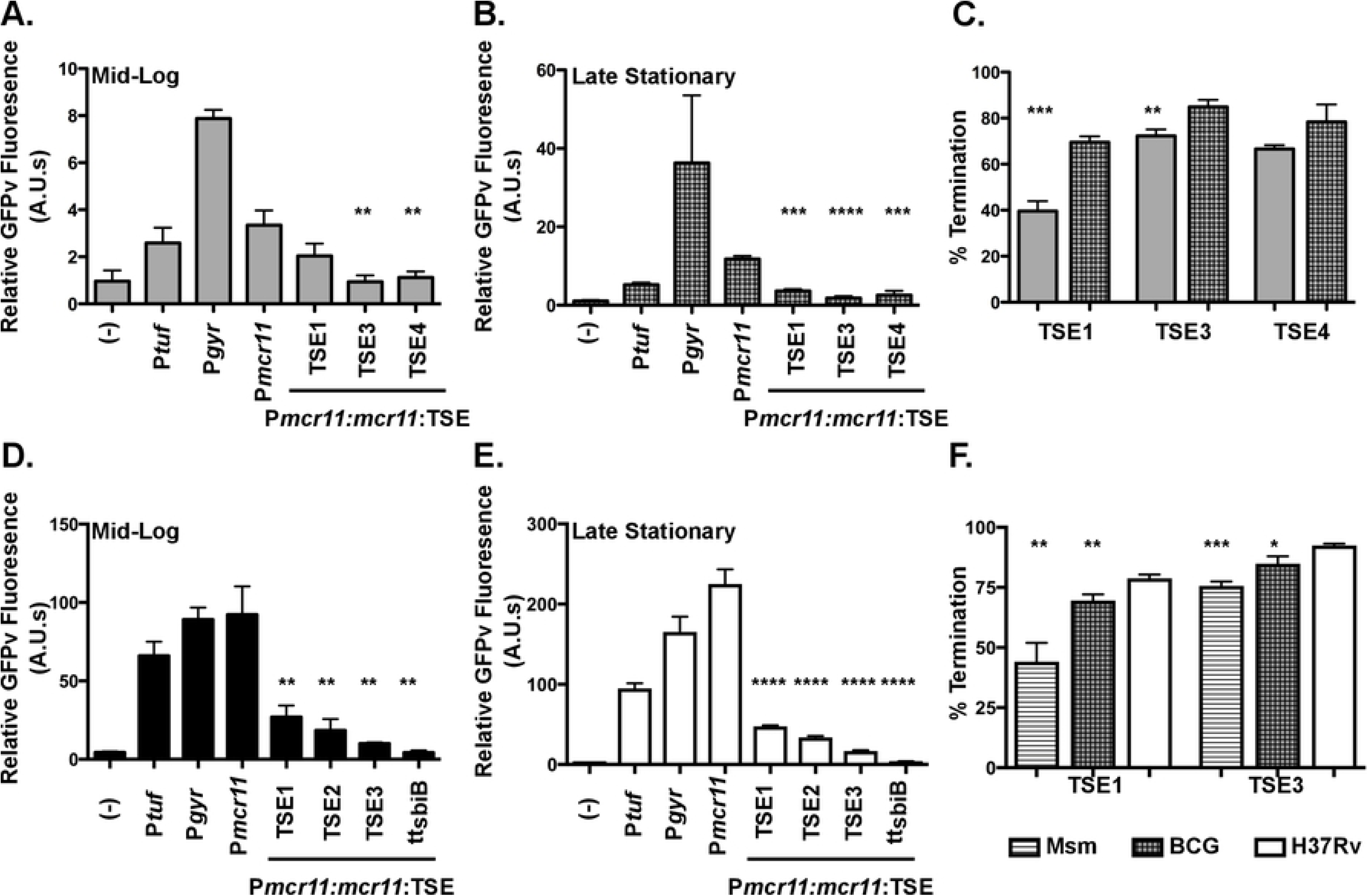
Efficiency of TSEs are positively regulated in response to growth phase in BCG and function significantly better in Mtb. A. GFPv fluorescence assay used to measure promoter activity and read-through of the TSEs of *mcr11* in mid-log phase BCG in hypoxic (1.3% O_2_, 5% CO_2_), shaking conditions. A promoterless (-) construct was used as a negative control and the promoters P*tuf* and P*gyr* served as positive controls. The various TSE constructs tested are indicated underneath the corresponding bar. Statistical comparisons were made to P*mcr11*. B. As in (A), but assayed in late stationary phase. C. A comparison of % termination observed in mid-log phase (solid bars) and late stationary phase (hatched bars) BCG. Statistical comparisons were made between mid-log and stationary phase. D. GFPv fluorescence assay used to measure promoter activity and read-through of the TSEs of *mcr11* in mid-log phase Mtb in hypoxic (1.3% O_2_, 5% CO_2_), shaking conditions. A promoterless (-) construct was used as a negative control and the promoters P*tuf* and P*gyr* serve as positive controls. The various TSE constructs tested are indicated underneath the corresponding bar. Statistical comparisons are made to P*mcr11*. E. As in (D), but assayed in late stationary phase. Statistical comparisons are made to P*mcr11*. F. A comparison of % termination observed in late stationary phase Msm, BCG, and Mtb. Statistical comparisons are made to Mtb. Results are the means of 3 biological replicates. Statistical analysis conducted with an unpaired, 2-tailed Student’s t-test. Asterisks indicate significance as follows: * p<0.05, ** p<0.01, *** p<0.001, **** p<0.0001.

### Longer TSEs enhance expression of Mcr11 in TB-complex mycobacteria

We expected that increased termination efficiencies would correlate with higher levels of mature Mcr11 in TB complex mycobacteria. Thus, we used Northern blot analyses to measure the production of stable Mcr11 from various *mcr11* TSE constructs in *Δmcr11* strains of BCG and Mtb during late stationary phase (Figure 4). Robust levels of Mcr11 were produced from constructs containing native *mcr11* TSEs. As was observed in experiments with Msm, Mcr11 size remained constant despite its expression with TSEs of different lengths in BCG or Mtb (Figure 4). This size restriction is consistent with precise termination or an RNA processing event that produces discrete 3’ ends.

**Figure 4.**
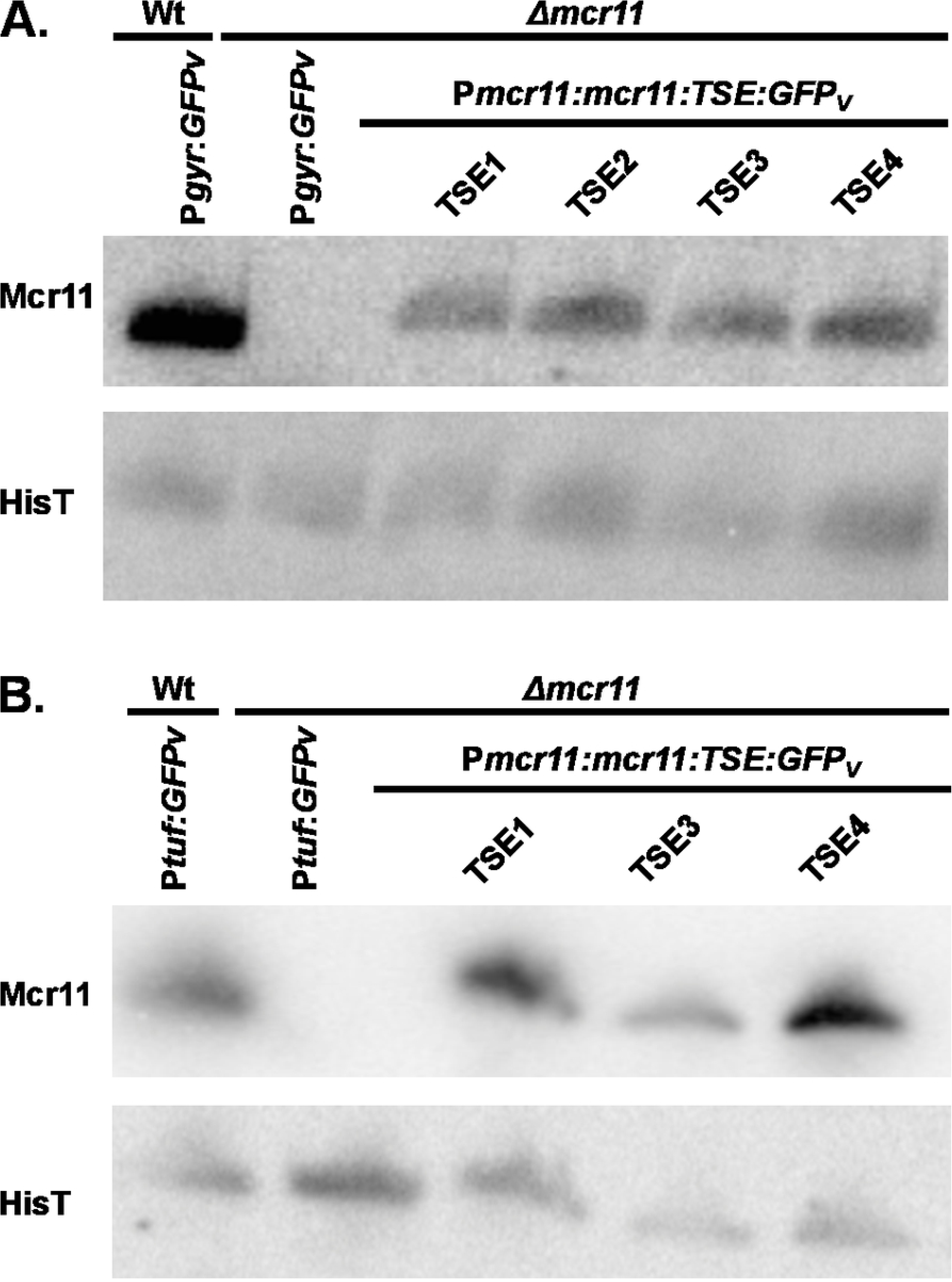
Different Mcr11 TSEs do not alter the size of stable Mcr11. A. Northern blot analysis of Mcr11 expression in the indicated strains of BCG in hypoxic (1.3% O_2_, 5% CO_2_) late stationary phase. B. Northern blot analysis of Mcr11 expression in the indicated strains of Mtb in hypoxic late stationary phase. The tRNA HisT is used as a loading control. Results representative of 2-3 independent repeats.

We considered the possibility that expression of *mcr11* in the context of an mRNA with a translated, stable transgene such as *GFPv* could contribute to the stability of Mcr11. Production of Mcr11 from *mcr11*:TSE1 constructs was compared to Mcr11 levels from a similar construct in which *GPFv* is not immediately downstream of TSE1 (Figure 5A). Mcr11 levels were not affected by the GFPv location (Figure 5B), indicating that *GFPv* did not contribute to the production or processing of stable Mcr11 in these experiments. This conclusion is also consistent with the low levels of Mcr11 observed in Msm despite the presence of the *GFPv* transgene (Figure 2D).

**Figure 5.**
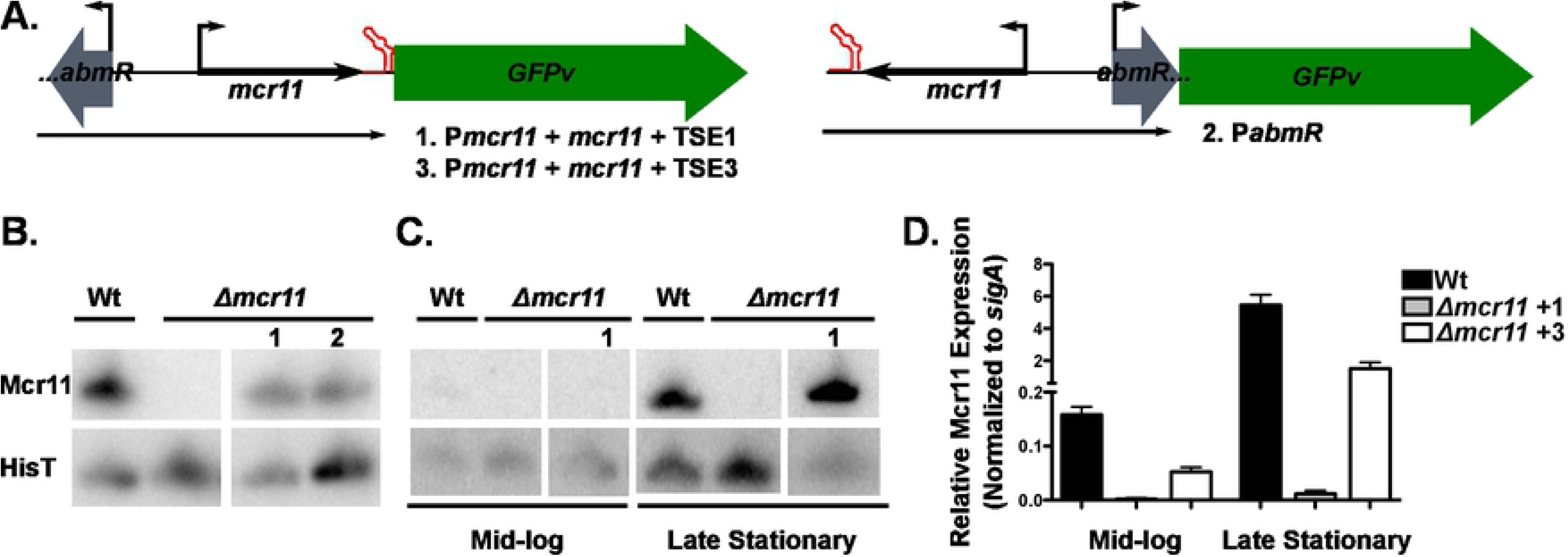
Termination at a sub-optimal TSE does not require a trans-gene for stabilization of Mcr11. A. A schematic of the transgene fusion constructs. *Mcr11* complementation was provided with (1) or without (2) the transgene GFPv fused to the end of *mcr11* downstream of sub-optimal terminator TSE1. An Mcr11 complementation construct was also tested with TSE3 (3). B. Northern blot analysis of Mcr11 expression in Mtb grown to late stationary phase in hypoxia (1.3% O_2_, 5% CO_2_). HisT was used as a loading control. C. Northern blot analysis Mcr11 expression across growth phase in BCG, indicated below. HisT was used as a loading control. D. qPCR analysis of Mcr11 expression in mid-log and late stationary phase BCG, normalized to *sigA* expression. Results are the representative of 2-3 biological replicates. Statistical analysis conducted with an unpaired, 2-tailed Student’s t-test. Asterisks indicate significance as follows: ** p<0.01, *** p<0.001.

As advancing growth phase significantly improved the termination efficiencies of TSEs in BCG (Figure 3C), we tested the effects of growth phase on the expression of *mcr11*. Mcr11 levels were observed by Northern blot (Figure 5C) and measured by qRT-PCR (Figure 5D) in BCG*Δmcr11* strains carrying ectopic copies of either *mcr11*:TSE1 or *mcr11*:TSE3. Both constructs showed growth-phase dependent increases in expression of stable Mcr11 (Figure 5C and 5D). Stress conditions can modulate the efficiency of RITs (47) and induce the appearance of multiple, different size products of a single sRNA in Mtb (23, 27). However, neither BCG nor Mtb displayed significant stress-induced changes in the termination efficiencies of *mcr11* TSEs (Supplemental Table 3) when subjected to nitrosative stress (DETA-NO), ATP depletion (BDQ), DNA damage (OFX), or transcriptional stress (RIF). These data demonstrate the significant role that differences in native 3’ sequence have on stable Mcr11 expression.

### Identification of Mcr11 regulatory targets

Having identified the sequence features required for robust ectopic production of Mcr11 in complemented *Δmcr11* strains, we considered *mcr11* function. A bioinformatic search of potential regulatory targets of Mcr11 using TargetRNA (48) and TargetRNA2 (49) identified *abmR* as the top-scoring hit (Figure 6A) (Supplemental Table 4). We measured the relative abundance of AbmR protein by Western blot analysis, and found that it was decreased in BGC*Δmcr11* and Mtb*Δmcr11* (Figure 6B,C). However, recent reports indicate that the *abmR* mRNA is a leaderless transcript that lacks a 5’ UTR (50), so the region of direct base-pair complementarity with the Mcr11 sRNA and the 5’ end of *abmR* predicted by TargetRNA is not likely to be present in the *abmR* mRNA (50). We also previously showed that the *mcr11* gene overlaps a substantial portion of the DNA sequences identified as the upstream enhancer and promoter regions for *abmR* transcription (Figure 6A) (33, 51).

**Figure 6.**
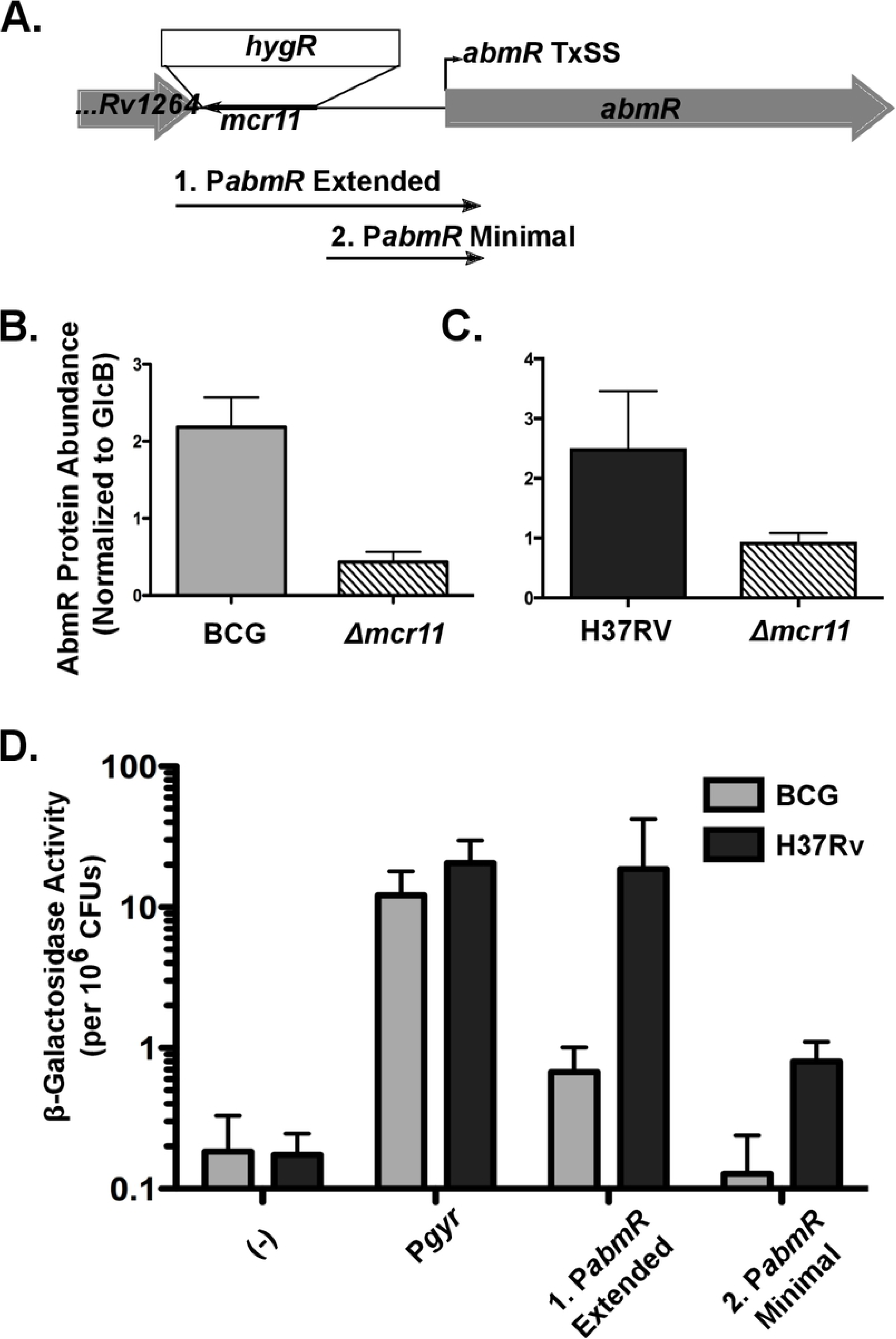
*Δmcr11* has a disrupted *abmR* promoter, resulting in reduced expression of AbmR protein in BCG and Mtb. A. The *mcr11* locus with the position of the hygromycin knockout cassette is shown. The regions of DNA used to create *promoter:lacZ* fusions are shown. Fragment (1.) includes the *Rv1264-abmR* intergenic region, which includes the *mcr11* locus. Fragment (2.) includes the *mcr11-Rv1265* sequence that is available in the *Δmcr11* strain. B. Quantification of western blot analysis from BCG grown to late-log phase in ambient, shaking conditions. C. Quantification of western blot analysis from Mtb grown to late-log phase in hypoxic (1.3% O_2_, 5% CO_2_), shaking conditions. AbmR was detected with poly-clonal anti-sera, and levels were normalized to GlcB levels, as detected by a monoclonal antibody. D. β-Galactosidase activity assays using the promoter:*lacZ* fusions shown in (A.) in Wt BCG (gray bars) and MTB (black bars) grown to late-log phase in ambient, shaking conditions. An asterisk * indicates p-value < 0.05 using an unpaired Student’s t-test. Results representative of 2-3 biological repeats.

We addressed the possibility that *mcr11* deletion has cis rather than trans effects on *abmR* expression due to disruption of *abmR* regulatory sequences. Promoter:*lacZ* reporter studies were used to compare P*abmR* activity in the presence and absence of flanking *mcr11* sequences in Wt BCG and Mtb backgrounds that express Wt levels of Mcr11. Transcriptional activity from the minimal *abmR* promoter lacking adjacent sequence in the *mcr11* gene was much lower than that from the extended *abmR* promoter that includes contiguous *mcr11* gene sequences (Figure 6D). These assays were conducted in the Wt backgrounds of BCG and Mtb with comparable levels of native *mcr11* expression in all matched strains, indicating that sequence overlapping the *mcr11* gene was required in cis for full *abmR* promoter activity. We conclude that the reduced expression of *abmR* in *Δmcr11* strains is likely due to the loss of sequence-based *abmR* regulatory elements within the *mcr11* gene rather than a trans-acting regulatory effect of Mcr11 on *abmR* expression.

TargetRNA also predicted multiple putative targets of Mcr11 regulation, including two genes within operons involved in central metabolic processes and cell division: *lipB* (encodes lipoate protein ligase B, needed for lipoate biosynthesis) (Figure 7A) and *Rv3282* (encodes a conserved hypothetical protein with homology to Maf septum-site inhibition protein) (Figure 7B). The intergenic spacing between putative target gene *fadA3* (*Rv1074c)*, which encodes a beta-ketoacyl coenzyme A thiolase, and the preceding gene (*Rv1075c*) is identical to that of the intergenic spacing between *Rv2216* and *lipB*. However, *fadA3* is likely to be transcribed independently of *Rv1075c* (50). *lipB* and Rv3282 were selected for follow-up because they are essential, associated with growth and central metabolism of Mtb, and contain predicted Mcr11 base-pairing sequences that are within known mRNA transcript boundaries. Each of these target transcripts has the potential to interact with Mcr11 through a 9-11 nt continuous base pairing region, followed by a shorter stretch of gapped or imperfect base pairing (Figure 7A and 7B). A region from nt 39-55 of Mcr11 was predicted to interact with these mRNA targets and others (Figure 7C and 7D, Supplemental Table 4), which includes a region of Mcr11 predicted to be in an unpaired loop in multiple modeled secondary structures of Mcr11 (Supplemental Figure 4).

**Figure 7.**
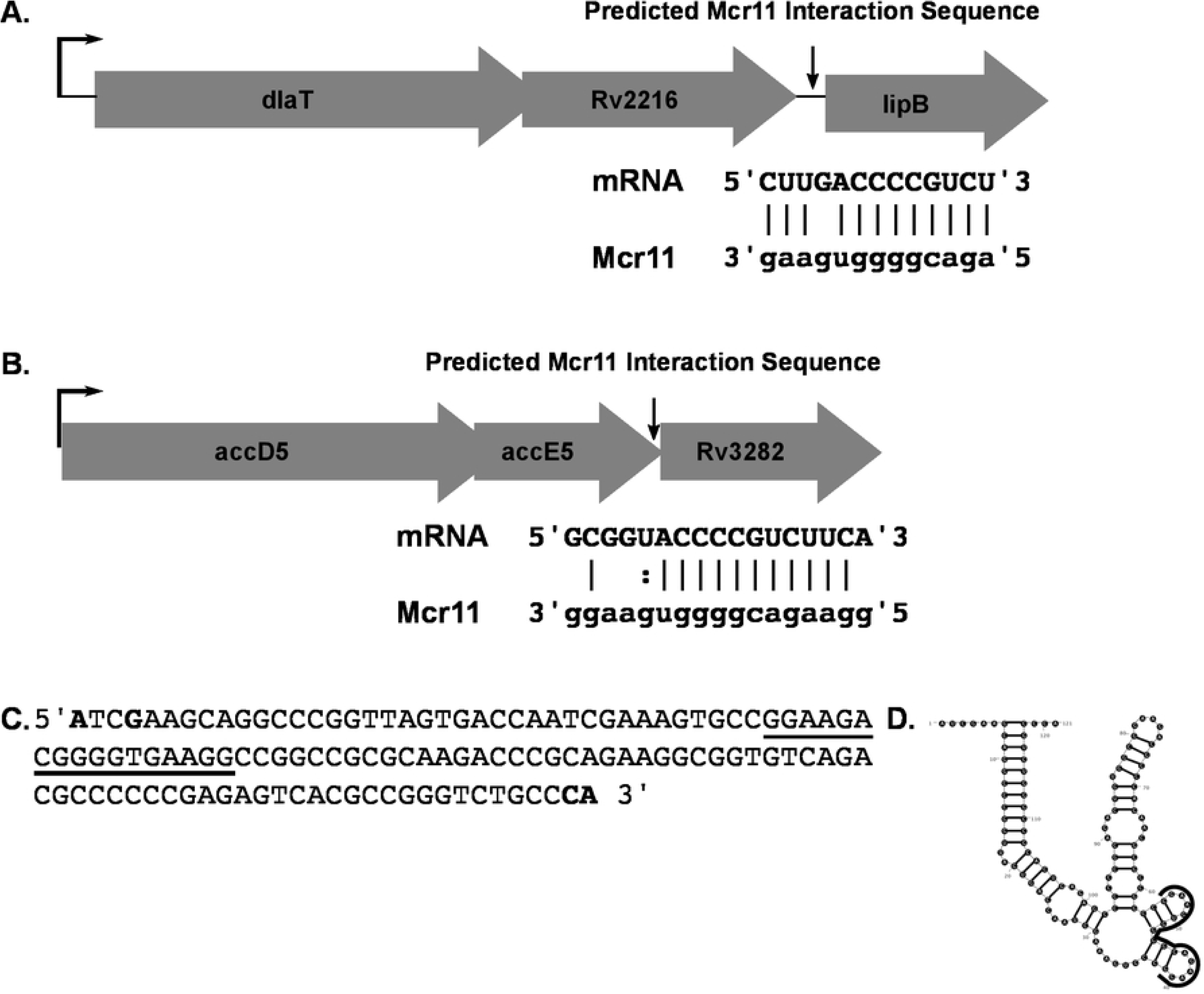
Bioinformatic modeling of Mcr11 targets reveals potential regulatory targets that are involved in central metabolism and cell division. A. The organization of the *dlaT-lipB* locus, with the position and potential base-pairing interactions between the mRNA and Mcr11 shown below. The transcriptional start site of the operon is shown with a thin black arrow. B. As in (A.), but for the *accD5-Rv3282* locus. Dashes indicate Watson-Crick base pairs, dots indicate non-Watson-Crick base pairs, and a blank space indicates no interaction between bases. C. The sequence of the Mcr11 RNA with mapped 5’ and 3’ ends shown in bolded letters. The portion of the sRNA predicted to interact with targets shown in (A) and (B) is underlined in black. D. The MFE secondary structure of Mcr11 as predicted by RNAStructure, with the portion of the sRNA predicted to interact with targets shown in (A) and (B) outlined in black.

### Fatty acids affect Mcr11-dependent regulation of target genes in Mtb

The *lipB* operon includes *dlaT*, which encodes DlaT, the E2 component of pyruvate dehydrogenase (PDH) and the peroxynitrite reductase/peroxidase (PNR/P) complex in Mtb. The function of DlaT is modulated by lipoylation, which is dependent upon lipoate biosynthesis by LipB (18, 52). BkdC (also called PdhC or Rv2495c) is a component of the branched chain keto-acid dehydrogenase (BCKADH) complex in Mtb that also requires lipoylation for activity. Disruption of components in any of these complexes can cause growth defects in Mtb, some of which are dependent upon the nutrient mixture present in the growth media (18, 19). The genes upstream of Rv3282 are *accD5* and *accE5*, which are needed for a long chain aceyl-CoA carboxylase enzymatic complex that generates substrates for fatty acid and mycolic acid biosynthesis (53, 54).

We used qRT-PCR to measure expression of genes within the operons containing predicted Mcr11-regulatory targets *lipB* and Rv3282 in Wt versus *Δmcr11* strains. Gene expression was queried in stationary phase BCG and Mtb grown in media with (+OA) or without (-OA) fatty acids to assess the impact of Mcr11 regulatory function in these conditions. Tween-80® is a hydrolysable detergent that can release a substantial amount of fatty acids (primarily oleic acid), and so was replaced with the non-hydrolysable detergent Tyloxapol in media lacking fatty acids. The *mcr11* complements included TSE4 with or without an intact copy of *abmR*. Levels of *phoP*, a response regulator expected to be independent of *mcr11* and *abmR*, and *pknA*, a serine-threonine protein kinase required for growth, were also measured.

Expression of *Rv3282*, Rv2216, *lipB* and *pknA* was significantly de-repressed in Mtb*Δmcr11* compared to Wt Mtb in the absence, but not the presence, of fatty acids. In contrast, levels of *lipB* and *pknA* expression decreased in the *mcr11* Mtb mutant when fatty acids were present. Complementation of *mcr11* fully (Rv3282, *pknA*) or partially (Rv2216, *lipB*) restored expression to Wt levels (Figure 8A). The expression of *accD5* (Rv3280) was significantly de-repressed and *accE5* (Rv3281) trended toward de-repression in Mtb*Δmcr11*, and complementation partially restored Wt levels of expression of both genes (Supplemental Figure 5A). Expression of *dlaT* and *phoP* were not altered in Mtb (Figure 8A). From these data, we concluded that Mcr11-mediated regulation expression of *Rv3282*, *Rv2216*, *lipB* and *pknA* expression in stationary phase Mtb is affected by the levels of fatty acids in the media. In contrast, no Mcr11-dependent regulation of the putative target genes was observed in BCG other than a trend of higher *Rv3282* expression in BCG*Δmcr11* relative to Wt BCG (Figure 8C and 8D). These data demonstrate that *mcr11*-dependent regulation of specific target genes is responsive to the fatty acid content of the culture media, and suggest that Mcr11 has a role in regulating the central metabolism of Mtb.

**Figure 8.**
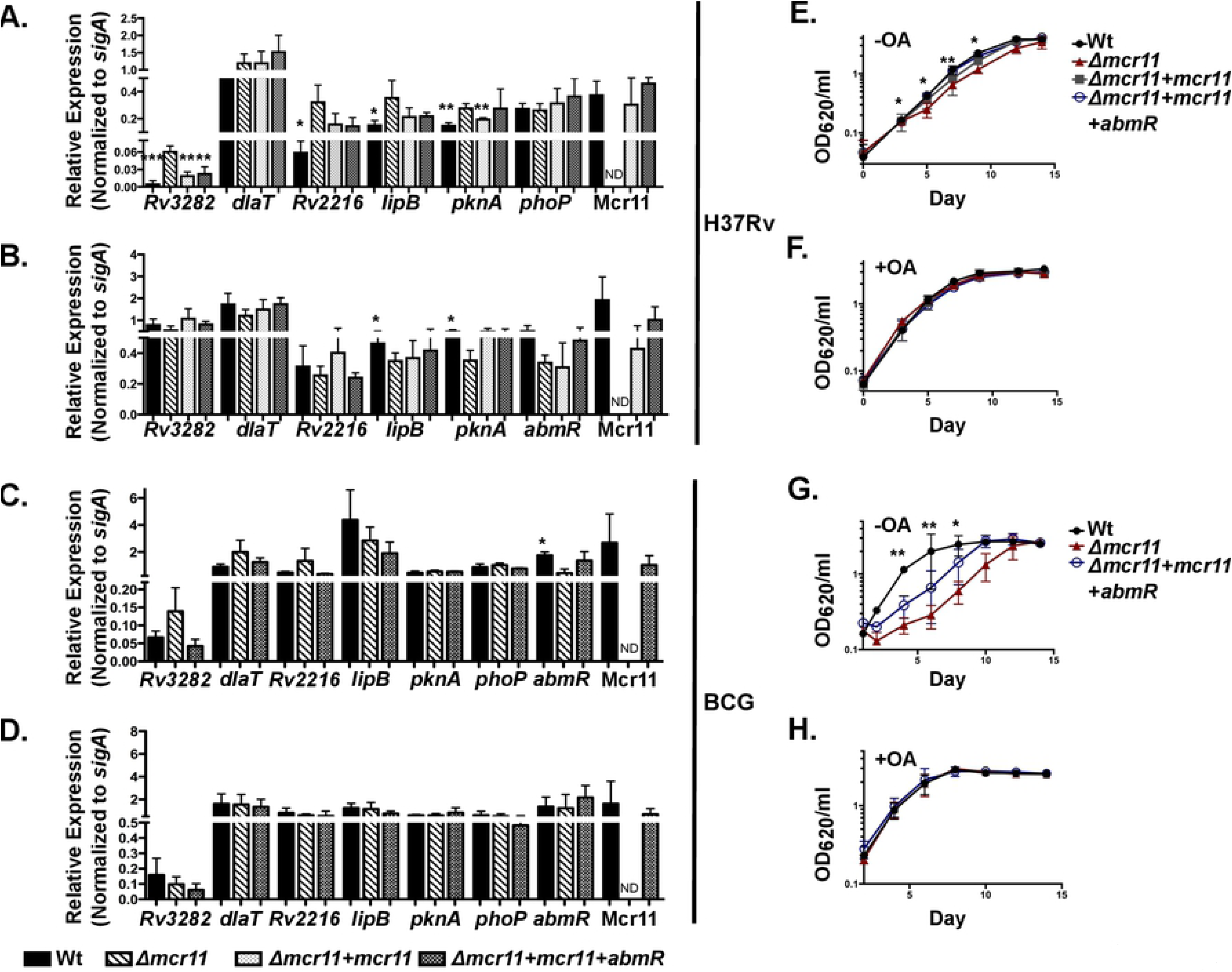
*Δmcr11* strains of BCG and Mtb are defective for growth in fatty-acid depleted media and predicted regulatory targets of Mcr11 are dysregulated at the mRNA level. A. Mtb was grown for 12 days in under hypoxic (1.3% O_2_, 5% CO_2_), shaking conditions in -OA media (7H9 + 0.2% glycerol, 10% ADC, and 0.05% Tyloxapol). Gene expression was measured by qRT-PCR and normalized to the reference gene *sigA*. Comparison made of each strain versus *Δmcr11*. Complementation strains included a single-copy of *mcr11* with TSE3 fused to *GFPv* or a single-copy of the *abmR* locus and *mcr11* with TSE4. B. Mtb was grown for 7 days in under hypoxic, shaking conditions in +OA media (7H9 + 0.2% glycerol, 10% OADC, and 0.05% Tyloxapol). Gene expression was measured by qRT-PCR and normalized to the reference gene *sigA*. C. BCG was grown for 12 days in under hypoxic, shaking conditions in -OA media. Gene expression was measured by qRT-PCR and normalized to the reference gene *sigA*. D. BCG was grown for 12 days in under hypoxic, shaking conditions in +OA media. Gene expression was measured by qRT-PCR and normalized to the reference gene *sigA*. E. Growth curve of Mtb grown in hypoxic, shaking conditions in -OA media. Growth was surveyed by measuring the optical density at 620 nm (OD_620_). F. Growth curve of Mtb grown in hypoxic, shaking conditions in +OA media. Growth was surveyed by measuring OD_620_. G. As in (E.), but with BCG. H. As in (F.), but with BCG. Results are the means of 3 biological replicates. Statistical comparisons made of each strain versus *Δmcr11* using an unpaired, 2-tailed Student’s t-test. Asterisks indicate significance as follows: * p<0.05, ** p<0.01, *** p<0.001, **** p<0.0001.

Expression levels of mRNA do not always match expression levels of the cognate protein if a gene is subjected to multiple layers of regulation. We tested if the protein levels of PknA or lipoylated DlaT were altered in Mtb*Δmcr11* by Western blot. No significant differences in the abundance of PknA or lipoylated DlaT were observed between Mtb*Δmcr11* and Wt Mtb (Supplemental Figure 5B). It is possible that de-repression of *lipB* does not affect DlaT lipoylation if lipoate is limiting or if wild-type levels are already saturating for lipoylation of DlaT.

### Mcr11 is required for optimal growth of BCG and Mtb without fatty acids

Optimal growth of Mtb on carbohydrate-based carbon sources requires appropriate *dlaT* expression (18, 19) and *lipB, accD5,* and *accE5* are essential for the growth of Mtb (55, 56). Mutations in *Rv3282* delay Mtb growth, even in nutrient rich media (55). Based on our observed regulation of these genes by Mcr11, we hypothesized that *Δmcr11* deleted strains would exhibit a growth defect when forced to utilize carbohydrate carbon sources for growth in media lacking a source of fatty acids.

No growth differences were observed between Wt and *Δmcr11* mutant Mtb or BCG in media containing fatty acids. However, BCG*Δmcr11* was severely growth-lagged and Mtb*Δmcr11* was moderately growth delayed compared to Wt bacteria when grown in media lacking fatty acids (Figure 8E-H). Complementation with *mcr11* partially complemented growth in BCG and fully restored growth to Wt levels in Mtb (Figure 8G and 8E). From these data, we conclude that *mcr11* has a role in the central metabolism and growth of BCG and Mtb.

## Discussion

This work identifies several genes associated with central metabolism in Mtb as regulatory targets of the sRNA Mcr11. We also demonstrated that transcriptional termination and stable production of the sRNA Mcr11 is enhanced by extended native sequence 3’ to *mcr11* along with the product of the divergently transcribed adjacent gene, *abmR*. The regulation of central metabolism in Mtb is consistent with *mcr11* expression profiles in response to growth phase and *in vivo* infection (26, 33). However, the additional requirements for TB complex specific factors for stable Mcr11 expression were unexpected, and this may have broader implications for understanding sRNA expression in Mtb.

Characterization of the factors required for efficient termination and stable expression of Mcr11 was a prerequisite for the complementation studies that confirmed Mcr11 regulatory targets. Protein coding mRNAs often can tolerate variable amounts of 5’ and 3’ flanking sequence because the signals for protein expression are provided immediately upstream and within the open reading frame (ORF). In contrast, expression of functional sRNAs may be more dependent on RNA chaperones and/or processing factors, as well as cis-acting sequence elements at their transcriptional boundaries that may be difficult to define (25, 47, 57-59). Defining the role of these cis-acting sequences for *mcr11* and possibly other sRNAs in Mtb is an important topic for future studies.

### Mechanism of mcr11 transcriptional termination

Sequences at the 3’ end of sRNAs can be critical for stability, function and interaction with RNA chaperones, although such chaperones have yet to be identified in mycobacteria (25, 59-64). The role of extended native 3’ sequences for expression of mycobacterial sRNAs warrants further investigation to determine whether *mcr11* is exceptional or representative of a larger group of sRNAs with regards to expression and stability requirements.

A recent study of Rho function in Mtb (41) found that depletion of Rho did not impact transcriptional boundaries at predicted RITs (43, 44), demonstrating a clear separation in the populations of transcripts terminated by RITs and Rho-dependent mechanisms. While we found that TSEs promote expression of Mcr11, their underlying mechanism remains unclear and our data do not precisely fit either the current RIT or Rho-dependent termination models (Table 1). We propose a model in which *mcr11* processing occurs immediately following, or concurrent with, Rho-dependent termination.

**Table 1:**
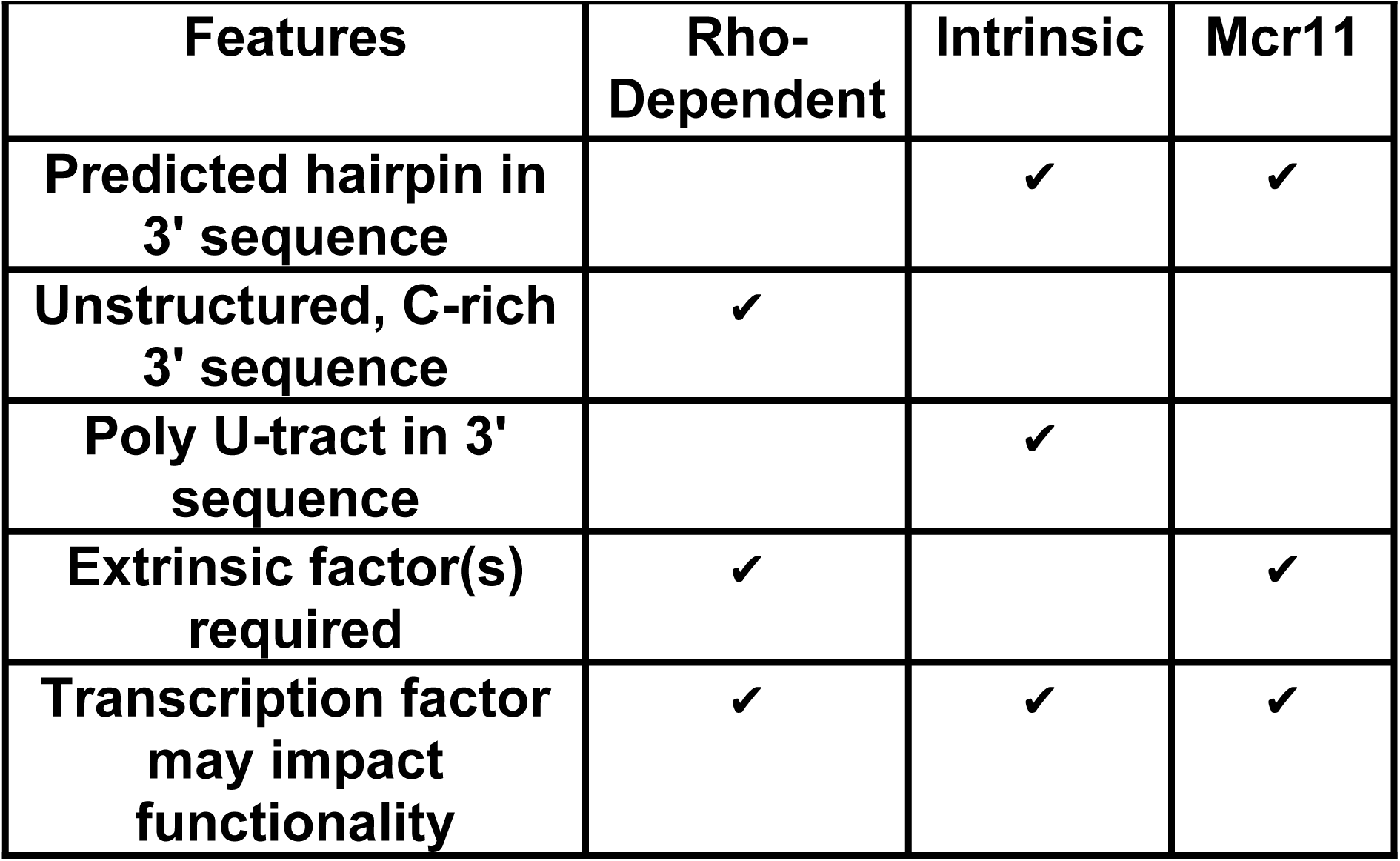
Comparison of transcription termination types

Rho-independent termination is considered “intrinsic” because extrinsic accessory factors are not required for function (38). Our results clearly indicate that production of stable Mcr11 requires factors other than the TSEs (Figs. 2D and 4). The species-specific effects we observed for *mcr11* termination measured by the GFPv reporter assays suggest that bacterial-specific factors affect the efficiency of *mcr11* termination itself, although the effects on Mcr11 stability are more striking (Figs. 2-4).

Rho utilization (rut) sites are degenerate and difficult to predict using bioinformatics approaches, but transcripts terminated by Rho tend to have C-enriched, G-depleted 3’ ends that are thought to be unstructured (40). It is not clear if there is a rut site within *mcr11*, as the sequences downstream of the 3’ end of Mcr11 are predicted to be highly structured and are rich in both G and C (Fig. 1). A run of 6Cs (nucleotides 93-98) occurs within Mcr11, but this region may be highly structured (Fig. 1). In contrast, Mcr11 TSEs 2-4, which contributed strongly to the termination of Mcr11, all possess C-rich loops in their predicted secondary structures that could be rut sites (Fig. 1). Rho-significant regions (RSRs) reported in the Botella et al study (41) include an RSR that begins 6 nt downstream of the mapped 3’ end of Mcr11, that is characterized as a region of general antisense transcription. The presence of this RSR is consistent with the possibility that *mcr11* expression is Rho-terminated. Mcr10 (*ncRv1157a*) is also proximal to an RSR (41). Future work is needed to establish the importance of Rho-dependent termination for mycobacterial sRNAs.

In *Escherichia coli*, multiple trans-acting factors are known to modulate termination and anti-termination of Rho-terminated sRNAs (65). AbmR, an ATP-responsive DNA-binding transcription factor that activates *mcr11* expression (33), was found here to also positively regulate termination efficiency of *mcr11*. Despite the high similarity between *abmR* and its Msm orthologue (33), *MSM_5010* failed to strongly activate *mcr11* expression and Mcr11 transcripts were unstable in Msm, even when *abmR* was provided in trans. It is possible that AbmR interacts directly with RNAP or Rho to affect termination, or recruits an as yet undefined trans-acting factor that modulates Mcr11 termination in Mtb. It will also be important to determine which specific sequence or structural elements in the TSEs of *mcr11* contribute to transcriptional termination versus processing, and the extent to which similar TSEs are associated with other sRNAs in mycobacteria.

The discrete 3’ end of mature Mcr11 observed by Northern blot is consistent with a processed or precisely terminated RNA, as transcription through TSEs would otherwise result in RNAs significantly longer products. Processing of the 3’ ends of mycobacterial sRNAs has been proposed (31), and a recent report identified a hypoxia-regulated mycobacterial sRNA that is extensively processed at its 3’ end (25). The observed size variants of specific sRNA species in response to host-associated stress conditions (23) provide further evidence for processing of sRNAs in mycobacteria.

Processing mechanisms for tRNAs and sRNAs have been defined, particularly for transcripts terminated by RITs in *E. coli* (57, 66-70). Recent work has demonstrated that Rho terminated transcripts have processed 3’ ends immediately downstream of a stable stem-loop (71). The processing of these discrete ends is dependent on the redundant action of the 3’ → 5’ exonucleases PNPase and RNase II, and it was speculated that the stem-loop promotes stabilization of the processed transcript (71). While few RNA processing enzymes of mycobacteria have been well characterized, the Mtb genome encodes many known RNases with varying sequence specificities (72-77). We speculate that unidentified TB-complex RNA chaperones and/or modifying enzymes contribute to Mcr11 stability, and that their absence in Msm results in Mcr11 degradation.

### Regulation of predicted targets of Mcr11

Oleic acid is the main fatty acid present in rich mycobacterial media formulations, provided either directly as oleic acid or in the hydrolyzable, nonionic detergent Tween-80®. The presence of Tween-80® can alter acid resistance (78), enhances the growth of mycobacteria when combined with glucose or glycerol (79, 80), and increases the uptake of glucose by BCG (79, 80). While we did not observe sensitivity to acid in BCG*Δmcr11* or BCG*ΔabmR* (data not shown), *mcr11* was required for growth on media lacking added fatty acids. Nutrient rich media is often used in batch culture experimentation with Mtb, and the potential confounding effect of multiple nutrient sources on characterizing the essentiality and function of gene products has recently gained appreciation (36, 81). Future studies characterizing nutrient uptake in Mtb and BCG strains of *Δmcr11* or *ΔabmR* will further our understanding of the nutrient related growth defects of these strains.

Two metabolic enzymes critical for Mtb pathogenesis are known to be lipoylated in Mtb, including dihydrolipoamide acyltransferase (DlaT) and BkdC, a component of the Lpd-dependent branched chain keto-acid dehydrogenase (BCKADH) (18). Lipoylation in Mtb is presumed to depend solely on lipoate synthesis by the enzymes LipA and the Mcr11 target LipB, as scavenging and import pathways are apparently lacking (52). *Rv3282* has homology to Maf, a septum inhibition protein conserved across all domains of life (82, 83). It is in a putative operon with *accD5* and *accE5*, which encode a probable propionyl-coenzyme A carboxylase involved in the detoxification of propionate using the methylmalonyl pathway to produce methyl-branched virulence lipids (84). Despite *mcr11*-dependent regulation of *Rv3282* in the absence of fatty acids, Mtb*Δmcr11* did not have a filamentous cell morphology (data not shown) and the function of Rv3282 in Mtb has not been defined. The regulation of Mcr11’s targets was responsive to the fatty acid content of the growth media, and it will be important to determine if the observed growth defects of Mtb*Δmcr11* are due to the dysregulation of *lipB* and *Rv3282*, or if there are additional targets of Mcr11 regulation that account for this phenotype (Figure 9). Additionally, the role of Mcr11 in supporting the growth and persistence of Mtb in response to nutrient availability and growth arrest should be explored.

**Figure 9.**
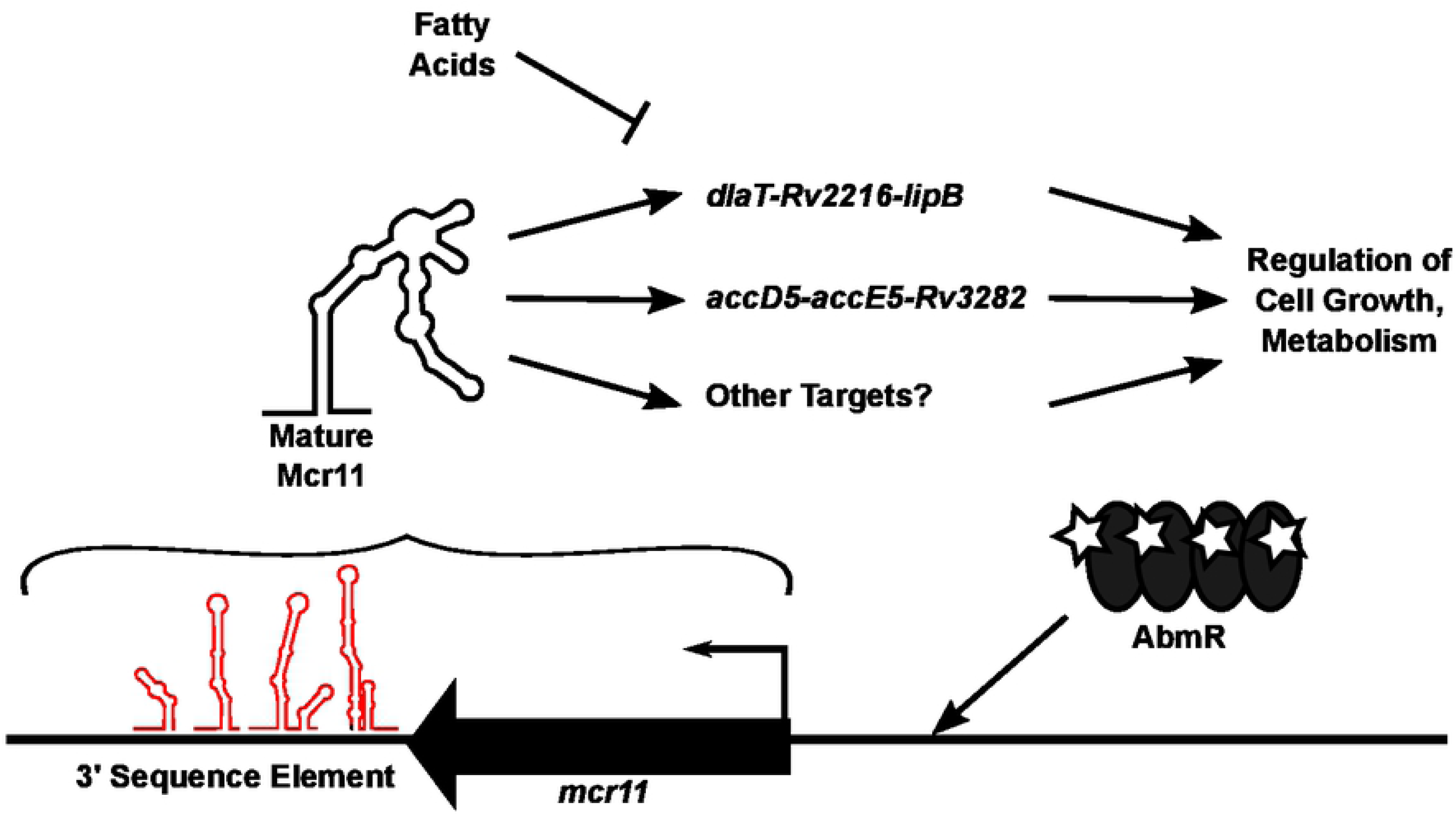
Model of Mcr11 function in Mtb. Expression of mcr11 is activated by cAMP, chronic infection in the lungs of mice, and by advancing growth phase and the ATP-binding transcription factor AbmR Native 3’ sequence elements (TSEs, predicted secondary structure shown in red) promote the transcriptional termination and stability of Mcr11. The predicted secondary structure of mature Mcr11 is shown in black. Mcr11 regulates the expression of genes involved in the central metabolism and growth of Mtb, possibly through a base-pairing interaction between Mcr11 and target mRNAs. Mcr11 is required for the optimal growth of Mtb in the absence of fatty acids

Mcr11-dependent regulation of *lipB* and *Rv3282* expression is expected to occur through base-paring with the mRNA, although future work is needed to establish the mechanism (Figure 7). The sRNA RyhB is known to differentially regulate expression of individual genes within the *iscRSUA* operon, resulting in down-regulation of the *icsSUA* genes while maintaining expression of *iscR* (85). This mechanism requires an intra-cistronic base-pairing dependent translational blockade of *iscS*, followed by recruitment of RNaseE and selective degradation of the 3’ end of the mRNA. The position of both putative Mcr11 base-pairing sites within target gene operons raises multiple possibilities for *mcr11*-mediated regulation of co-transcribed genes that will be an intriguing topic for future study. We noted that the *mcr11* target interaction site within the intergenic region of *Rv2216* and *lipB* had identical spacing to that of *Rv1075c* and *fadA3*, although the significance of this architecture is unknown.

This study shows that the transcriptional termination and stability of the sRNA Mcr11 is enhanced by extended, native 3’ sequence elements (TSEs) in TB complex mycobacteria. The role of AbmR in *mcr11* expression was extended from that of a transcriptional activator to include enhancement of *mcr11* transcriptional termination. Our observation that bacterial-species specific factors govern sRNA stability in mycobacteria may also extend to other sRNAs. Combined use of bioinformatic and molecular tools established fatty acid responsive, *mcr11*-dependent gene regulation in Mtb and provides a versatile strategy for the continued search of sRNA function in pathogenic mycobacteria. Future work defining the precise roles of the sRNA Mcr11 in regulating the growth and metabolism of Mtb will greatly advance our understanding of this important pathogen.

## Materials and Methods

### Bacterial Strains and Growth Conditions

*Mycobacterium tuberculosis* H37Rv (Mtb) (ATCC 25618) and *Mycobacterium bovis* BCG (BCG) (Pasteur strain, Trudeau Institute) were grown on 7H10 agar (Difco) supplemented with 10% oleic acid-albumin-dextrose-catalase (OADC) (Becton Dickson and Company) and 0.01% cyclohexamide or in Middlebrook 7H9 liquid medium (Difco) supplemented with 10% (vol/vol) OADC, 0.2% (vol/vol) glycerol, and 0.05% (vol/vol) Tween-80 (Sigma-Aldrich) in physiologically relevant shaking, hypoxic conditions (1.3% O_2_, 5% CO_2_) (86) for promoter::reporter fusion assays and Northern blot analysis. *Mycobacterium smegmatis* mc^2^155 (Msm) (ATCC 700048) was grown in Middlebrook 7H9 liquid medium (Difco) supplemented with 10% (vol/vol) OADC, 0.2% (vol/vol) glycerol, and 0.05% (vol/vol) Tween-80 (Sigma-Aldrich), shaking in ambient air for promoter::reporter fusion assays and Northern blot analysis. All experiments were started from low-passage frozen stocks. For cloning experiments, *Escherichia coli* strains were grown on Luria-Bertani (Difco) agar plates or in liquid broth. Growth media and plates were supplemented with 50 μg/mL hygromycin, 25 μg/mL kanamycin (Sigma-Aldrich) for selection of clones. All bacterial cultures were incubated at 37°C.

### Mutant Strain Construction

Knockout strains of *mcr11* in Mtb and BCG genetic backgrounds were generated using homologous recombination (87) to replace nucleotides 1-100 with a hygromycin resistance cassette. Disruption of *mcr11* was confirmed with polymerase chain reaction (PCR) (Supplemental Figure 1B. and 1C.) and Northern blot analysis (Supplemental Figure 1D. and 1E.). Several complementation constructs were created to restore Mcr11 expression. Sequence from the corrected *abmR* (*Rv1265)* start site (51) to the end of the *Rv1264* ORF was amplified by PCR and cloned downstream of the *efTu* (Rv0685) promoter in the multi-copy plasmid pMBC280, creating pMBC1211. PCR was used to amplify varying lengths of sequence from the end of the *Rv1264* ORF through the end of the *abmR* ORF, and they were cloned it into pMBC409, which contains a kanamycin resistance marker, to create pMBC2040, pMBC2041, and pMBC2042 (Supplemental Figure 1A.). Complementation plasmids were sequenced to verify that the desired sequences had been correctly incorporated and then used to transform Mtb and BCG by electroporation. A complete list of plasmids used in this study are provided in Supplemental Table 1 and primers are listed in Supplemental Table 2.

### Mapping the 3’ End of Mcr11

RNA was harvested from late-log phase Mtb and the 3’ end of Mcr11 was mapped using the miScript Reverse Transcription Kit (Qiagen) (88) and a gene-specific primer. The reaction product was cloned into pCRΙΙ-TOPO vector (Life Technologies) and ten positive clones were subjected to Sanger sequencing. The sequences were mapped back to the chromosome of Mtb.

### Bioinformatic Modeling of Mcr11

Secondary structure modeling of Mcr11 and putative native intrinsic terminators was performed using RNAstructure using default parameters (46). Color-coded base-pair drawings of predicted RNA secondary structures were created with Visualization Applet for RNA (VARNA) (89) and simplified line drawings were made in Inkscape (inkscape.org). A synthetic terminator (tt_sbi_B) (90) was modeled onto Mcr11 as well. Secondary structure modeling of Mcr11 was performed using RNAstructure (46). Using the previously published 5’ and 3’ boundaries of Mcr11, minimum free energy (MFE) and centroid structures for Mcr11 derived from CentroidFold (using the CONTRAfold inference engine), Mfold, NUPACK, RNAfold, and RNAStructure algorithims were created (46, 91-94). Base-pair drawings of predicted RNA secondary structures were either directly exported from the software used, or created with Visualization Applet for RNA (VARNA) (89) and simplified line drawings were made in Inkscape (inkscape.org).

Regulatory targets of Mcr11 in Mtb were predicted using TargetRNA (48) and TargetRNA2 algorithms (49), set to default parameters (Table3). Where possible, the position of predicted regions of base-paring between Mcr11 and a putative regulatory target were evaluated against known 5’ ends of mRNAs in Mtb (50) (Table 3).

### Promoter::Reporter Fusion Assays

Promoter:l*acZ* fusions were created to compare the relative transcriptional activities from DNA sequences spanning the intergenic space between *Rv1264* and *abmR*, and *mcr11* and *abmR* (Supplemental Figure 2). The promoters of *gyrB* (*Rv0005*) and *efTu* (*Rv0685*) were used as positive controls, and a promoterless construct was used as a negative control. The relevant constructs were transformed into wild-type (Wt) strains of BCG and Mtb and assayed for β-galactosidase activity in late log phase cultures grown in ambient, shaking conditions by adding 5-acetylaminofluoresceindi-β-D-galactopyranoside (C_2_FDG; Molecular Probes) and measuring fluorescence in a Cytofluor 4000 fluorometer (PerSeptive Biosystems) as described previously (95).

Constructs including the *mcr11* promoter, Mcr11 sequence, and various potential intrinsic terminator sequences were fused to the reporter green fluorescent protein Venus (*GFPv*) on an integrating plasmid as previously described (33). A GFPv fluorescence assay was used to measure promoter activity and read-through of the putative intrinsic terminators of Mcr11 in Msm, BCG, and Mtb.

At mid-log phase and late stationary phase, aliquots of recombinant strains were collected and gently sonicated (setting 4, 4 pulses of 5” on time interspersed by 5’ off time) using a Virsonic 475 Ultrasonic Cell Disrupter with a cup horn attachment (VirTis Company) before duplicate samples were diluted 1:1 in fresh media. The level of fluorescence in arbitrary units from the GFPv reporter strain was detected using the CytoFluor Multi-Well Plate Reader Series 4000 (PerSeptive Biosystems) at 485 nm excitation and 530 nm emission. The optical density (OD) at 620 nm was read using a Tecan Sunrise® microplate reader and fluorescence levels were normalized to 10^6^ bacteria. A promotorless (-) construct was used as a negative control and the promoters P*tuf* and P*gyr* served as positive controls. The % termination was calculated by dividing the observed GFPv fluorescence in arbitrary units of each indicated terminator construct by the amount of GFPv fluorescence in arbitrary units measured from the *mcr11* promoter.

Late stationary phase reporter strains of BCG and Mtb were diluted with fresh media to an OD620 nm of 1.25 in a 12 well plate, and then subjected to a variety of cellular stressors for a 24 hour period in shaking, hypoxic conditions (1.3% O_2_, 5% CO_2_). Cells were treated with 250 μM diethyltriamine-NO (DETA-NO) (Sigma-Aldrich), 1 μg per ml ofloxacin (OFX), 0.28 μg per ml rifampacin (RIF), 53 μg per ml bedaquiline (BDQ), or 0.1% dimethyl sulfoxide (DMSO) vehicle control. GFPv fluorescence in arbitrary units per 106 bacteria (by measured OD at 620 nm) was monitored as described above, and percent termination was calculated relative to P*mcr11* controls.

### Fluorescence-Activated Cell Sorting Analysis

Fluorescence-activated cell sorting analysis was performed on Msm samples that had been fixed with 4% paraformaldehyde (PFA) in phosphate buffered saline (PBS) at pH 7.0 for 30 minutes at room temperature. Fixation was quenched pH-adjusted glycine, and cells were washed three times with PBS. Cells were diluted approximately 1:500 in DPBS and subjected to FACS analysis with a FACS Calibur (Becton Dickson) as previously described (96). Data was collected for 20,000 events per sample and analyzed in CellQuest software (Becton Dickenson).

### RNA Isolation

Total RNA was harvested from Msm, BCG and Mtb after cultures were treated 0.5 M final GTC solution (5.0 M guanadinium isothiocyanate, 25 mM sodium citrate, 0.5% sarkosyl, and 0.1 M 2-β-mercaptoethanol) and pelleted at 4°C. Pelleted cells were resuspended in TRIzol Reagent (Invitrogen) and cells were disrupted with 0.1 mm zirconia-silica beads (BioSpec Products) and three 70-second high-speed pulse treatments in a bead-beater (BioSpec Products). RNA was recovered from lysates with Direct-zol Mini Prep columns (Zymo) per the manufacturer’s protocol. RNA was eluted from the column with nuclease free-water, treated with DNaseI (Qiagen) for 20 minutes at room temperature, and isopropanol precipitated. One μg of total RNA was screened for DNA contamination by PCR using primers KM1309 and KM1310 internal to the sigA ORF. The quality of each RNA sample was assessed by running 1.0 μg of DNA-free RNA on a 0.1% (weight/vol) SDS-agarose gel and visualizing intact 23S, 16S, and 5S ribosomal RNA bands after staining with ethidium bromide (Sigma-Aldrich).

### Northern Blots

High quality DNA-free total RNA isolated from Msm, BCG, and Mtb was used for Northern blot analysis of Mcr11 expression. 3-10 μg RNA was separated on a 10% 8 M urea PAGE run at a constant current of 20 mA for 1-1.5 h. The gel was electroblotted onto a Hybond N (Millipore) nylon membrane using a wet transfer system (Bio-Rad Laboratories) as previously described (Girardin, in prep). Blots were UV cross-linked and baked at 80°C for 2 h prior to pre-hybridization at 42°C for 1 h in Rabid-Hyb Buffer (GE Healthcare Life Sciences). Hybridization with α-^32^P-ATP end-labeled DNA oligo probes was performed at 42°C for 2-16 h. Blots were washed per the manufacturer’s protocol and exposed to phosphor-screens for visualization.

### Generation of Mycobacterial Cell Lysates and Western Blotting

Strains of BCG and Mtb were grown in the desired condition, and pelleted and washed twice in ice-cold Dulbecco’s phosphate buffered saline, calcium and magnesium free (DPBS-CMF), with 0.2% protease inhibitor (Sigma-Aldrich). Cell pellets were resuspended in 1/25 volume lysis buffer (0.3% SDS, 200 mM DTT, 28 mM Tris-Hcl, 22 mM Tris-Base, and 1% protease inhibitor cocktail) and disrupted by two rounds of high-powered sonication at 4°C with Virsonic 475 Ultrasonic Cell Disrupter with a cup horn attachment (VirTis Company) interspersed with 10 freeze-thaw cycles as previously described (33). Lysate was cleared by centrifugation and the cleared lystate was heat-killed at 95°C before quantification with a NanoDrop 2000 (Thermo Scientific).

A total of 30 μg protein of each sample was separated by 12-15% Tris-glycine SDS-PAGE, and gels were immunoblotted on Immobilon-P membranes (Millipore) for 1h at 1 mA/cm^2^ using a wet transfer system (Bio-Rad Laboratories). Blots were blocked and proved with 1° antibody in 5% milk (vol/vol) in 50 mM Tris-buffered saline with 0.05% (vol/vol) Tween-20 (Fisher Scientific). Monoclonal antibodies against *Mycobacterium tuberculosis* GlcB (Gene Rv1837c), Clone α-GlcB (produced *in vitro* NR-13799) was obtained through the NIH Biodefense and Emerging Infections Research Resources Repository, NIAID, NIH. Polyclonal mouse AbmR anti-serum was previously generated in-house (33). Polyclonal rabbit anti-lipoic acid antibody (EMD Millipore catalogue # 437695-100ul) was obtained from Fisher Scientific. Primary antibodies were detected with peroxidase conjugated goat secondary antibody and enhanced chemiluminescence (ECL) western blotting detection reagent (Thermo Scientific).

### Growth Curve Analysis

The growth of BCG and Mtb in hypoxic (1.3% O2, 5% CO2), shaking conditions were monitored by measuring the optical density at 620 nm (OD620) of gently sonicated aliquots of cells with a Tecan Sunrise® microplate reader. Cultures were grown in vented T25 tissue culture flasks (Corning), and time points were in single or multi-day intervals. Growth comparisons between Middlebrook 7H9 + 0.2% glycerol, 10% OADC, and 0.05% Tween-80 (+OA) or in Middlebrook 7H9 + 0.2% glycerol, 10% ADC, and 0.05% Tyloxapol (-OA) were made.

## Acknowledgements

We are very grateful to Dr. Guangchun Bai and Damen Schaak for generating the *mcr11* knockout strains of BCG and Mtb used in this study. We also thank Dr. Joseph Wade for helpful discussions, and gratefully acknowledge the Wadsworth Center Applied Genomics Technologies Core for DNA sequencing and the Wadsworth Center Media and Tissue Culture Core for media preparation.

## Supporting information

**S1 Figure. Generation and confirmation of *mcr11* knock-out and complementation strains.** A. A diagram of the *mcr11* locus and the relative position of the hygromycin resistance cassette. Lines with arrows indicated the DNA sequence included in the various listed complementation strains. B. PCR confirmation of *mcr11* knockout and complementation strains of BCG. C. PCR confirmation of *mcr11* knockout and complementation strains of Mtb. D. Northern blot analysis of the indicated strains of BCG from 3 μg of total RNA harvested from late stationary phase cultures grown in hypoxic (1.3% O_2_, 5% CO_2_), shaking conditions. E. Northern blot analysis of the indicated strains of Mtb from 5 μg of total RNA harvested from late-log phase cultures grown in hypoxic (1.3% O_2_, 5% CO_2_), shaking conditions. The number above the lane indicates the identity of each sample. The column to the left of the rows indicates which gene was amplified or which RNA was probed by Northern blot.

**S2 Figure. FACS analysis of TSEs in Msm.** FACS analysis of GFPv fluorescence of 20,000 events for each sample as indicated by the label to the left of the histogram. Representative of three independent repeats.

**S3 Figure. *abmR* contributes to the termination of Mcr11 transcripts.** A. GFPv fluorescence assay used to measure m*cr11* promoter activity in mid-log phase Mtb in hypoxic (1.3% O_2_, 5% CO_2_), shaking conditions. B. The % termination in mid-log phase. The various TSE constructs tested are indicated underneath the corresponding bar. C. Promoter reporter assay as in (A), but in late stationary phase. D. The % termination, as in (B), but in late stationary phase. Results representative of 3 biological replicates. Statistical analysis conducted with an unpaired, 2-tailed Student’s t-test. Comparison made versus Mtb*ΔabmR* in A. and C., and versus Wt in B. and D. Asterisks indicate significance as follows: * p<0.05, ** p<0.01, *** p<0.001, **** p<0.0001.

**S4 Figure. Secondary structure of Mcr11 RNA generated by multiple modeling algorithms.** A. The CentroidFold secondary structure using the CONTRAfold inference engine. B. The −54.40 kcal/mol MFE structure generated using Mfold. C. The −45.40 kcal/mol MFE strcture generated using NUPACK. D. The RNAfold centroid structure. E. The −54.00 kcal/mol MFE structure generated using RNAfold. F. The −53.80 kcal/mol MFE structure generated using RNAStructure. G. The ensemble centroid structure generated using Sfold. H. The −54.40 kcal/mol MFE structure generated using Sfold. S5 Figure. Protein levels of predicted regulatory targets of Mcr11 are not dysregulated in Mtb*Δmcr11* during hypoxia in the absence of added fatty acids. A. Mtb was grown for 12 days in under hypoxic, shaking conditions in -OA media. Gene expression was measured by qRT-PCR and normalized to the reference gene *sigA*. Comparison made of each strain versus Mtb*Δmcr11*. B. Mtb was grown for 12 days in under hypoxic (1.3% O_2_, 5% CO_2_), shaking conditions in Middlebrook 7H9 + 0.2% glycerol, 10% ADC, and 0.05% Tyloxapol (-OA) media. Protein levels were assayed by Western blot analysis with polyclonal α-Lipoate, polyclonal α-PknA, and monoclonal α-GlcB antibodies. GlcB serves as a loading and transfer control. Representative of three biological repeats.

**S6 Figure. Regulation of predicted targets of Mcr11 is condition specific in BCG and Mtb.** A. BCG grown for 12 days in under hypoxic (1.3% O_2_, 5% CO_2_), shaking conditions in Middlebrook 7H9 + 0.2% glycerol, 10% OADC, and 0.05% Tyloxapol (+OA). Gene expression was measured by qRT-PCR and normalized to the reference gene sigA. G.-H. Mtb was grown for 7 days in under hypoxic (1.3% O_2_, 5% CO_2_), shaking conditions in Middlebrook 7H9 + 0.2% glycerol, 10% OADC, and 0.05% Tween (+OA) and treated with vehicle control (Control) or 10 mM dibutyryl cAMP on day 3 (+db cAMP). Gene expression was measured by qRT-PCR and normalized to the reference gene *sigA*. Representative of 2-3 biological repeats. Statistical analysis conducted with an unpaired, 2-tailed Student’s t-test. Comparison made of each strain versus *Δmcr11*. Asterisks indicate significance as follows: * p<0.05, ** p<0.01, *** p<0.001, **** p<0.0001.

S1 Table. Plasmids used in the study. S2 Table. Primers used in the study.

S3 Table. Stress does not impact the efficiency of TSEs in BCG or Mtb.

S4 Table. Results of TargetRNA and TargetRNA2 predictions of Mcr11 regulatory targets.

